# Understanding disease-associated metabolic changes in human colon epithelial cells using *i*ColonEpithelium metabolic reconstruction

**DOI:** 10.1101/2024.10.22.619644

**Authors:** Boyu Jiang, Nick Quinn-Bohmann, Christian Diener, Vignesh Bose Nathan, Yu Han-Hallett, Lavanya Reddivari, Sean M. Gibbons, Priyanka Baloni

## Abstract

The colon epithelium plays a key role in the host-microbiome interactions, allowing uptake of various nutrients and driving important metabolic processes. To unravel detailed metabolic activities in the human colon epithelium, our present study focuses on the generation of the first cell-type specific genome-scale metabolic model (GEM) of human colonic epithelial cells, named iColonEpithelium. GEMs are powerful tools for exploring reactions and metabolites at systems level and predicting the flux distributions at steady state. Our cell-type-specific iColonEpithelium metabolic reconstruction captures genes specifically expressed in the human colonic epithelial cells. The iColonEpithelium is also capable of performing metabolic tasks specific to the cell type. A unique transport reaction compartment has been included to allow simulation of metabolic interactions with the gut microbiome. We used iColonEpithelium to identify metabolic signatures associated with inflammatory bowel disease. We integrated single-cell RNA sequencing data from Crohn’s Diseases (CD) and ulcerative colitis (UC) samples with the iColonEpithelium metabolic network to predict metabolic signatures of colonocytes between CD and UC compared to healthy samples. We identified reactions in nucleotide interconversion, fatty acid synthesis and tryptophan metabolism were differentially regulated in CD and UC conditions, which were in accordance with experimental results. The iColonEpithelium metabolic network can be used to identify mechanisms at the cellular level, and our network has the potential to be integrated with gut microbiome models to explore the metabolic interactions between host and gut microbiota under various conditions.

**Graphical Abstract:** 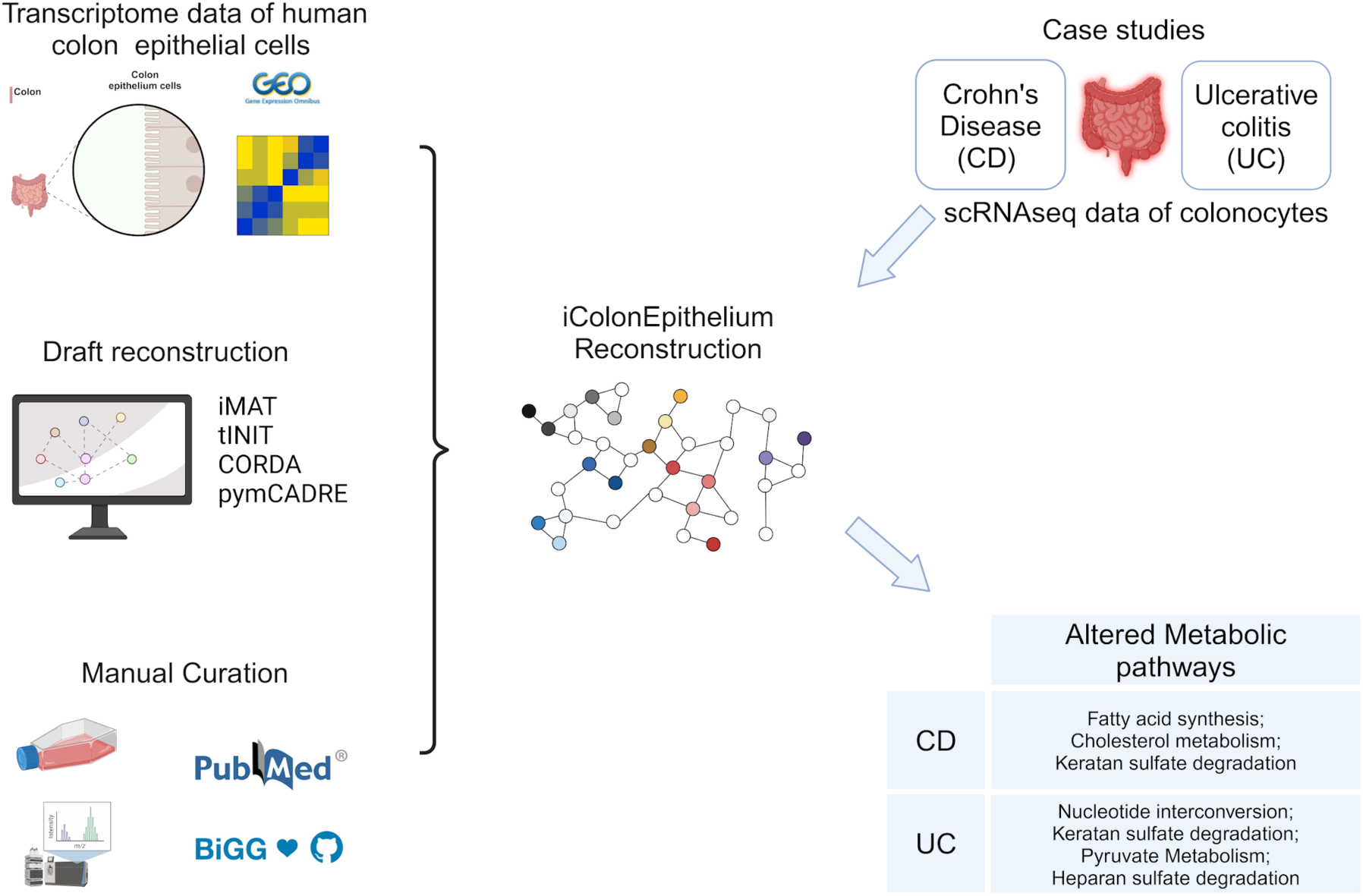

## Introduction

The human colon plays an important role in the host metabolism, as it not only contains the highest amount and density of microbiota in the human body, but also has the largest surface area that interacts with gut microbiota and coordinates nutrient absorption (Dupont et al., 2020). Homeostasis of the colon mainly depends on colonic epithelial cells, which form an epithelial barrier and regulate mucosal immunity during interactions with gut microbiota (Odenwald & Turner, 2017). Studies have indicated that a disrupted colon epithelial barrier is associated with intestinal diseases, including inflammatory bowel disease (IBD), systemic disorders such as type 1 diabetes, and neurological disorders (Groschwitz & Hogan, 2009; Martini et al., 2017; Pellegrini et al., 2023). These studies primarily focused on intestinal permeability and intestinal barrier integrity. Tight junction proteins expressed by colonocytes, including occludin, claudins, and zonula occludins, are important for normal barrier function and are regulated by many factors such as diet, pathogens, and environmental stress (Chelakkot et al., 2018).

In addition to participating in the composition of the physical barrier, recent studies have also highlighted the importance of metabolic functions in colonic epithelial cells. For example, the oxygen consumption of colonocytes, which contribute to mitochondrial β-oxidation of short-chain fatty acids (SCFAs), maintains anaerobic conditions in the gut and thus shapes the gut microbiome (Litvak et al., 2018a). Lactate produced from colonocytes can affect the composition of the gut microbiota during gut inflammation (Taylor et al., 2022). Metabolic circuits between colon epithelium and gut microbiome can potentially influence the blood metabolome (Diener et al., 2022; Rath & Haller, 2022). Apart from interacting with the gut microbiome, colonic epithelial cells can also affect immune cell functions through specific metabolic circuits. For instance, arginine and tryptophan, catabolized in colonic epithelial cells, were reported to have immunoregulatory effects in many diseases (Rath & Haller, 2022). Therefore, exploring the comprehensive and detailed metabolic network of colonocytes and their relationships with the gut microbiota is a promising avenue towards identifying microbiome-mediated therapeutic strategies.

Due to the complexity of the metabolic interactions between colon epithelium and gut microbiota, computational tools such as genome-scale metabolic modeling are required, which can improve both comprehensiveness and specificity of such interactions. Genome-scale metabolic models (GEMs) are network-based tools representing biochemical information in mathematical format. These *in silico* reconstructions can integrate transcriptome, metabolome, and other omics data (Fang et al., 2020; Passi et al., 2022) and enable us to simulate and predict metabolic fluxes through these reaction networks (Orth et al., 2010; Q. Wang et al., 2006). GEMs of the gut microbiome and the human tissues have been repeatedly applied in studies of host-microbiome interactions to facilitate the exploration of the mechanisms underlying metabolic disorders and the development of potential dietary interventions (Sen & Orešič, 2019; Thiele et al., 2020; van der Ark et al., 2017; Quinn-Bohmann et al., 2024).

To obtain more insights into the metabolic activities of the human colon epithelium, we present a cell-type-specific GEM of the human colonic epithelial cell, called iColonEpithelium. iColonEpithelium can achieve essential metabolic functions of the human colonic epithelial cell and enables exploration of the metabolic changes in the cells across diverse conditions. The reconstruction has been made context-or condition-specific by integration of transcriptome and metabolome data to study the underlying mechanisms involved in various health-related outcomes. By defining specific transport reactions of the co-metabolites between colonic epithelial cells and the gut microbiome, and adding them to iColonEpithelium, we have the ability to potentially simulate and predict interactions between the gut barrier and gut microbiome.

## Results

### Metabolic reconstruction of human colonic epithelial cells, iColonEpithelium

We generated the first human colonic epithelial cell-type-specific metabolic network using the generic human reconstruction, Recon3D (Brunk et al., 2018), as the template (see Methods). iColonEpithelium metabolic reconstruction has 6651 reactions, 4072 metabolites, and 1954 genes (Table S1a, S1b, S1c). We used transcriptome data of colonic epithelial cells from healthy individuals for generation of the reconstruction (Table S2). We obtained draft reconstructions using four established tools for generating context-specific metabolic reconstructions (see Methods section). Metabolites and reactions from all four drafts were compared and combined into a consensus reconstruction, that was finally refined for the generation of iColonEpithelium reconstruction. We compared iColonEpithelium reconstruction with the generic colon organ reconstruction, published as part of the human whole-body model (Thiele et al., 2020). Since our reconstruction was generated with transcriptomics data from colon epithelium, it primarily reflected metabolic features specific to colonocytes instead of the whole colon organ. For example, our reconstruction contained the reaction of mitochondrial acetate metabolism, which is catalyzed by acetyl-CoA synthetase. This reaction is not included in the colon part of the human whole-body model but can contribute to a better understanding of mitochondrial energy metabolism of colonocytes. Furthermore, objective functions of the iColonEpithelium were also decided based on the metabolic function of colonocytes, which put more emphasis on metabolization of short chain fatty acids (SCFAs). As shown in Figure 1, we found that approximately 37% and 95% of reactions in our reconstruction overlapped with reactions in the colon part of the whole-body model and Recon3D, respectively. The percentages of the overlapped metabolites and genes in the iColonEpithelium with the colon reconstruction are about 44% and 86%, respectively; while in comparison with Recon3D, our reconstruction shared 96% metabolites, and all its genes overlapped with Recon3D (details in Figure 1). Moreover, subsystems in the iColonEpithelium (Table S3) covered 103 metabolic pathways and the distribution of subsystem demonstrated that the top 10 most frequent subsystems belonged to transport reactions, fatty acid metabolism and lipid metabolism (Table 1), which suggests important metabolic functions of colonocytes involve exchanges and utilization of lipids, especially fatty acids (Rath & Haller, 2022; Schwärzler et al., 2024). Because previous studies have emphasized that beta-oxidation of short chain fatty acids (SCFAs) plays an important role in the physiological function of human colonic epithelial cells (Litvak et al., 2018a), we used biomass maintenance and SCFAs production as the objective functions in our reconstruction.

**Figure 1.**
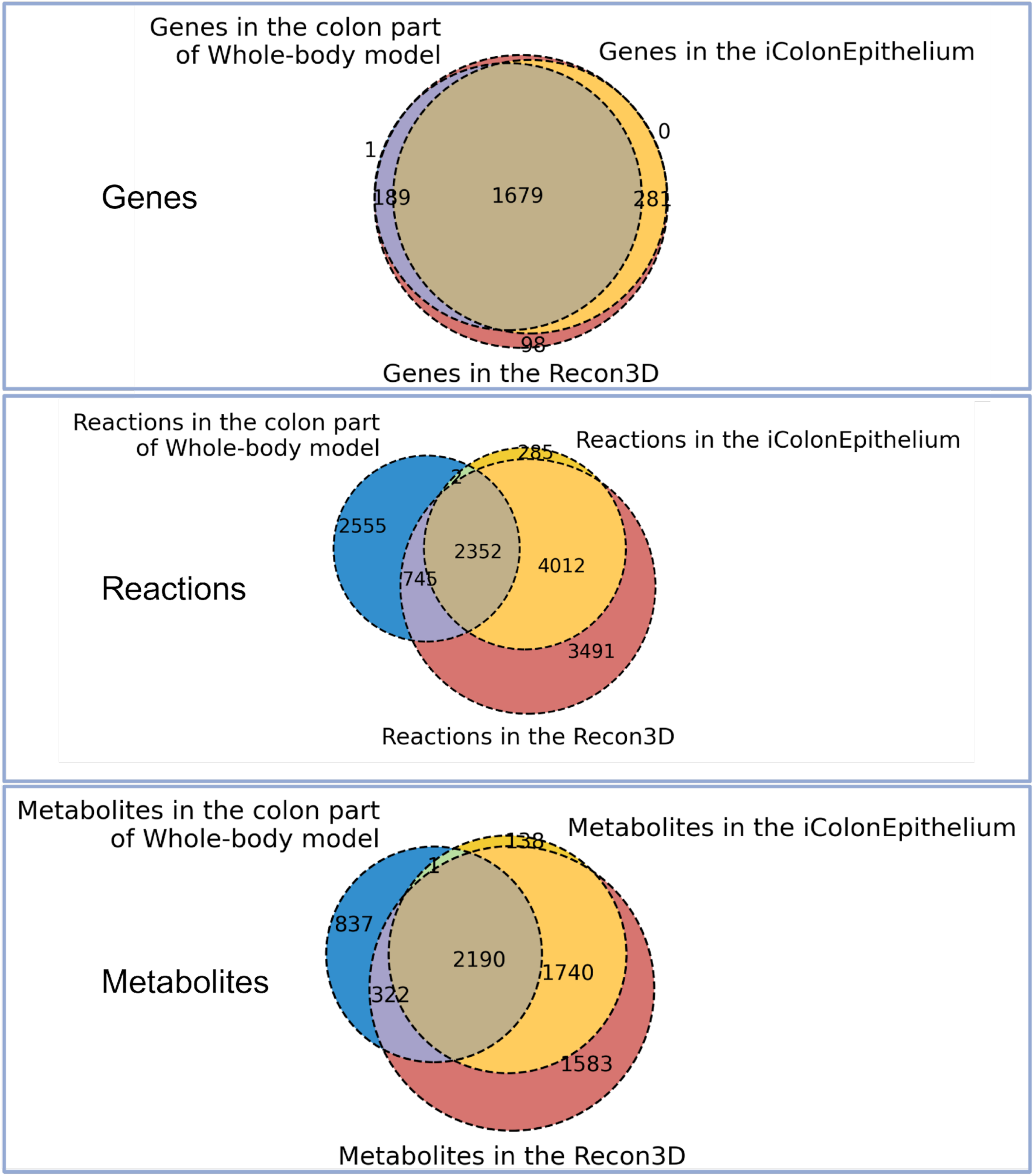
Venn diagrams indicating the overlap of genes, reactions and metabolites between iColonepithelium, Recon3D and colon part of Whole-body model (Harvey and Harvetta)

**Table 1.** Distribution of top 10 subsystems in the iColonEpithelium

| Subsystem | Frequency |
| --- | --- |
| Transport, extracellular | 1482 |
| Fatty acid oxidation | 857 |
| Extracellular exchange | 715 |
| Transport, mitochondrial | 275 |
| Cholesterol metabolism | 234 |
| Fatty acid synthesis | 228 |
| Glycerophospholipid metabolism | 140 |
| Nucleotide interconversion | 137 |
| Transport, lumen | 137 |
| Lumen exchange | 137 |

As a next step to test the specificity of our reconstruction, we tested iColonEpithelium reconstruction for human colonic epithelial-specific metabolic tasks. A total of 229 metabolic tasks associated with basic mammalian cells or human colonic epithelial cell functions were identified from published literature (Table S4). The consensus reconstruction passed ∼84% of them, including the β-oxidation of butyrate and acetate to generate ATP (Litvak et al., 2018a), and the synthesis of cholesterol and spermidine (Blachier et al., 2011; Phan & Tso, 2001), which were reported as important metabolic functions for colonocytes. These refinement steps suggested that the iColonEpithelium reconstruction is cell-type-specific and can be used for predicting the metabolic functions of the cell.

### Identification of transport reactions to enable connections between colonic epithelial cells and gut microbiota

To enable iColonEpithelium reconstruction to interact with the gut microbiota, we collected information regarding transport reactions and added them to the reconstruction. We used the information from the Human Protein Atlas (https://www.proteinatlas.org) (Uhlen et al., 2019) and 624 of these genes (Table S5) mapped to the reconstruction. We mined the information of the metabolic reactions corresponding to mapped genes from the Uniprot (https://www.uniprot.org) and Rhea (https://www.rhea-db.org) database (Bansal et al., 2022; Bateman et al., 2023), which provided us with 247 metabolites involved in the transport reactions. Information regarding transport reactions and corresponding metabolites from a published transport module (Sahoo et al., 2014) and the colon part of the whole-body model (Thiele et al., 2020) were also extracted and used for reconstruction. After comparing all metabolites and checking their transport in the EMBL database (https://www.embl.org) (Leinonen et al., 2011), we finalized on 139 metabolites that connect the reconstruction to the gut microbiota in the colon lumen, including SCFAs and bile acids (Table S6). We assigned these metabolites to the ‘lumen’ compartment and their corresponding transport reactions added to the reconstruction. With the lumen compartment and curated transport reactions, the iColonEpithelium reconstruction is able to simulate metabolic activities of colonic epithelial cells interacting with the gut microbiota.

### Metabolomic profiling and validation of iColonEpithelium using Caco-2 cell culture

We carried out *in vitro* culturing of colon carcinoma cell line (Caco-2) and performed targeted metabolomics to quantify concentrations of specific metabolites, such as amino acids and SCFAs in the cell culture medium. Since Caco-2 cells have been widely used as an *in vitro* model to study the function and metabolism of intestinal epithelium (Fedi et al., 2021), the changes of metabolite concentrations in the medium can reflect the capacity of colonocytes in utilizing or producing specific metabolites. We cultured Caco-2 cells with Dulbecco’s Modified Eagle Medium (DMEM) supplemented with fetal bovine serum (FBS) for 2 days and performed targeted metabolomics analysis of 20 amino acids. The cells mainly consumed 7 amino acids (glutamine, glutamate, isoleucine, leucine, phenylalanine, serine and valine) from the medium, while proline was secreted into the medium (Table 2). Previous reports have shown increased production of proline in Caco-2 cells (Alaqbi et al., 2022). The concentrations of butyrate and acetate in the cell culture medium decreased by 33% and 54%, respectively, compared with original medium whose concentrations of butyrate and acetate were 0.04 and 18.43 ng/μl, respectively (Table 2). It has been demonstrated that colonic epithelial cells could utilize butyrate and acetate from the extracellular environment, which is in accordance with reported studies (Stein et al., 2000), and the iColonEpithelium also contained necessary reactions which allowed uptake of butyrate and acetate for ATP generation. SCFAs were not specifically added to the DMEM, but their presence in the medium can be attributed to the fetal bovine serum (FBS), as it has been reported that FBS contains various fatty acid components (Lee et al., 2023).

**Table 2.**
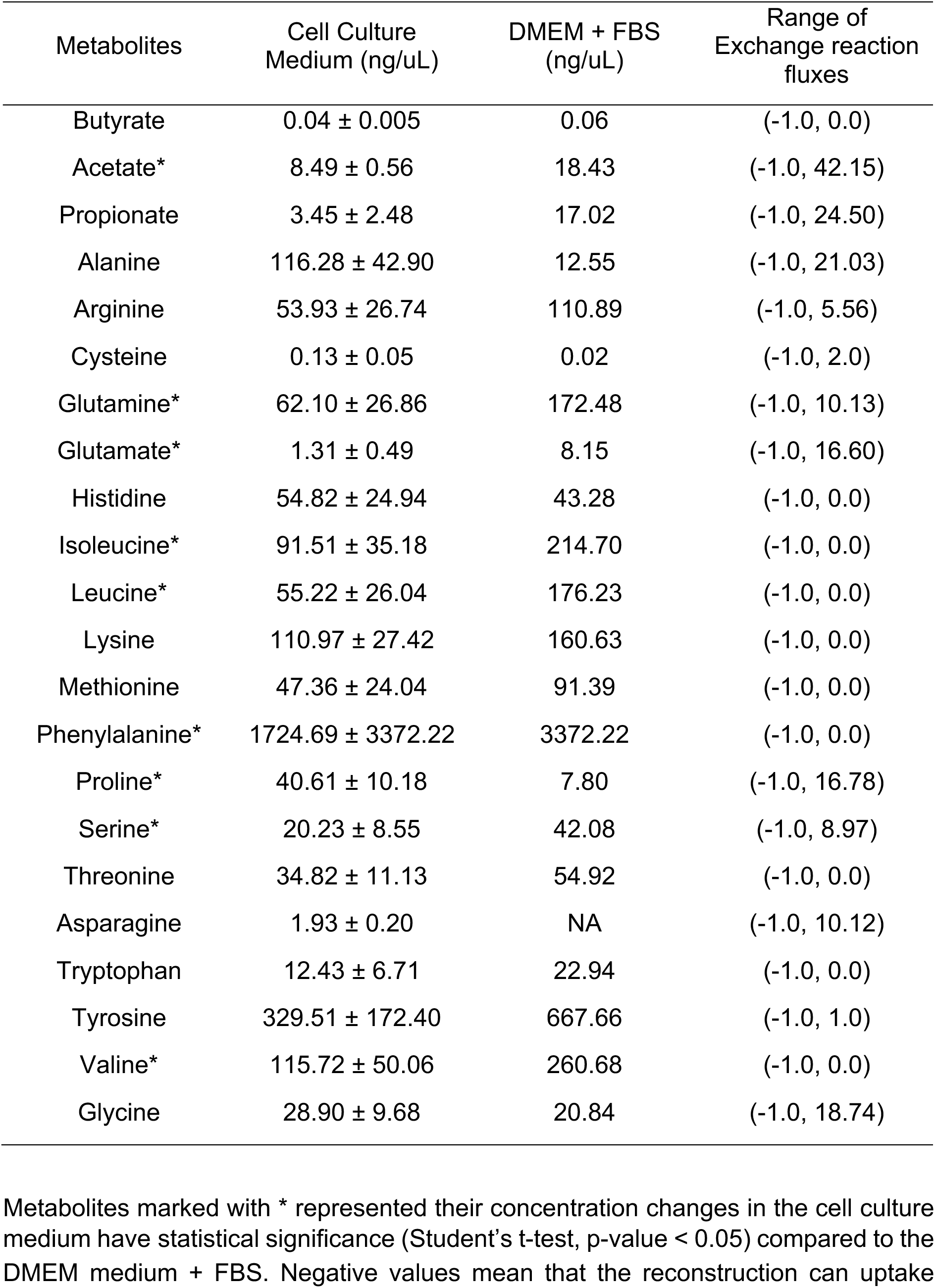
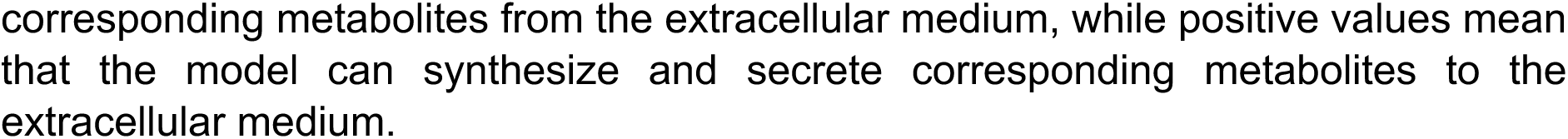
Targeted metabolomic analysis on metabolites from Caco-2 cells culturing medium

| Metabolites | Cell Culture Medium (ng/uL) | DMEM + FBS (ng/uL) | Range of Exchange reaction fluxes |
| --- | --- | --- | --- |
| Butyrate | 0.04 ± 0.005 | 0.06 | (-1.0, 0.0) |
| Acetate* | 8.49 ± 0.56 | 18.43 | (-1.0, 42.15) |
| Propionate | 3.45 ± 2.48 | 17.02 | (-1.0, 24.50) |
| Alanine | 116.28 ± 42.90 | 12.55 | (-1.0, 21.03) |
| Arginine | 53.93 ± 26.74 | 110.89 | (-1.0, 5.56) |
| Cysteine | 0.13 ± 0.05 | 0.02 | (-1.0, 2.0) |
| Glutamine* | 62.10 ± 26.86 | 172.48 | (-1.0, 10.13) |
| Glutamate* | 1.31 ± 0.49 | 8.15 | (-1.0, 16.60) |
| Histidine | 54.82 ± 24.94 | 43.28 | (-1.0, 0.0) |
| Isoleucine* | 91.51 ± 35.18 | 214.70 | (-1.0, 0.0) |
| Leucine* | 55.22 ± 26.04 | 176.23 | (-1.0, 0.0) |
| Lysine | 110.97 ± 27.42 | 160.63 | (-1.0, 0.0) |
| Methionine | 47.36 ± 24.04 | 91.39 | (-1.0, 0.0) |
| Phenylalanine* | 1724.69 ± 3372.22 | 3372.22 | (-1.0, 0.0) |
| Proline* | 40.61 ± 10.18 | 7.80 | (-1.0, 16.78) |
| Serine* | 20.23 ± 8.55 | 42.08 | (-1.0, 8.97) |
| Threonine | 34.82 ± 11.13 | 54.92 | (-1.0, 0.0) |
| Asparagine | 1.93 ± 0.20 | NA | (-1.0, 10.12) |
| Tryptophan | 12.43 ± 6.71 | 22.94 | (-1.0, 0.0) |
| Tyrosine | 329.51 ± 172.40 | 667.66 | (-1.0, 1.0) |
| Valine* | 115.72 ± 50.06 | 260.68 | (-1.0, 0.0) |
| Glycine | 28.90 ± 9.68 | 20.84 | (-1.0, 18.74) |

Metabolites marked with \* represented their concentration changes in the cell culture medium have statistical significance (Student's t-test, p-value < 0.05) compared to the DMEM medium + FBS. Negative values mean that the reconstruction can uptake
corresponding metabolites from the extracellular medium, while positive values mean that the model can synthesize and secrete corresponding metabolites to the extracellular medium.

To determine if the iColonEpithelium was able to predict the uptake or secretion of metabolites observed in the Caco-2 cell culturing experiment, we performed flux variability analysis (FVA) on the metabolic reconstruction. FVA in metabolic modeling is a computational technique used to determine the range of possible fluxes through reactions in GEM providing insights into flexibility of the network (Thiele et al., 2013). Based on our FVA results, we identified reactions, especially exchange reactions that are required by our reconstruction to import or secrete metabolites from or to the extracellular environment, like the culture medium, blood or the gut lumen. If the minimum flux of an exchange reaction in a GEM is negative value, this represents uptake from the extracellular environment, namely the medium, into the intracellular space; similarly, if the maximum flux of an exchange reaction in a GEM is positive value, this metabolite is secreted from the intracellular space to the extracellular environment. By simulating the uptake of metabolites from DMEM medium in our reconstruction (Table 2) and then running FVA, we found that the iColonEpithelium was able to uptake the 7 amino acids mentioned above, as well as synthesize proline with the DMEM, which is concurrent with our empirical observations.

The sanity checks (Brunk et al., 2018) on the iColonEpithelium confirmed that it passes the basic requirements, like containing no leaks in the reconstruction, generating no ATP from water, and maintaining flux consistency (Table S7). The reconstruction was also queried in the Memote tool (Lieven et al., 2020), which ensured the quality of iColonEpithelium. All the results of sanity check are provided in Supplementary File 2.

### Identifying the metabolic signatures in ulcerative colitis and Crohn’s disease using iColonEpithelium reconstruction

#### Generate context-specific reconstructions from single-cell RNA sequencing data

We integrated single-cell RNA sequencing (scRNAseq) data from patient samples with the iColonEpithelium reconstruction to investigate metabolic changes in two types of inflammatory bowel diseases (IBD): Crohn’s disease (CD) (Kanke et al., 2022) and ulcerative colitis (UC) (Smillie et al., 2019). The CD study consisted of transcriptome data of colonocytes from 4 healthy and 3 CD samples (Kanke et al., 2022) and we generated context-specific metabolic networks for each sample in this study using iMAT algorithm (Zur et al., 2010). Similarly for the UC study, we generated 36 context-specific reconstructions from 23 healthy samples and 13 UC samples (Smillie et al., 2019).

#### Understanding the metabolic differences in CD and UC cases

To explore metabolic differences in CD and UC conditions, we performed flux variability analysis on the context-specific metabolic reconstructions. We calculated minimum and maximum fluxes of reactions in each context-specific reconstruction. We compared the maximum reaction fluxes between the healthy and IBD samples in the CD and UC studies and carried out statistical analysis to identify reactions with significant flux differences. In the CD study, we identified 12 reactions whose maximum fluxes are significantly different with respect to the healthy controls. These reactions were mainly associated with fatty acid metabolism, keratan sulfate degradation and creatine uptake (Table 3a). In the UC study we identified 22 reactions with significant flux differences between healthy and UC groups that were part of nucleotide metabolism, and metabolite transportation (Table 3b).

**Table 3.**
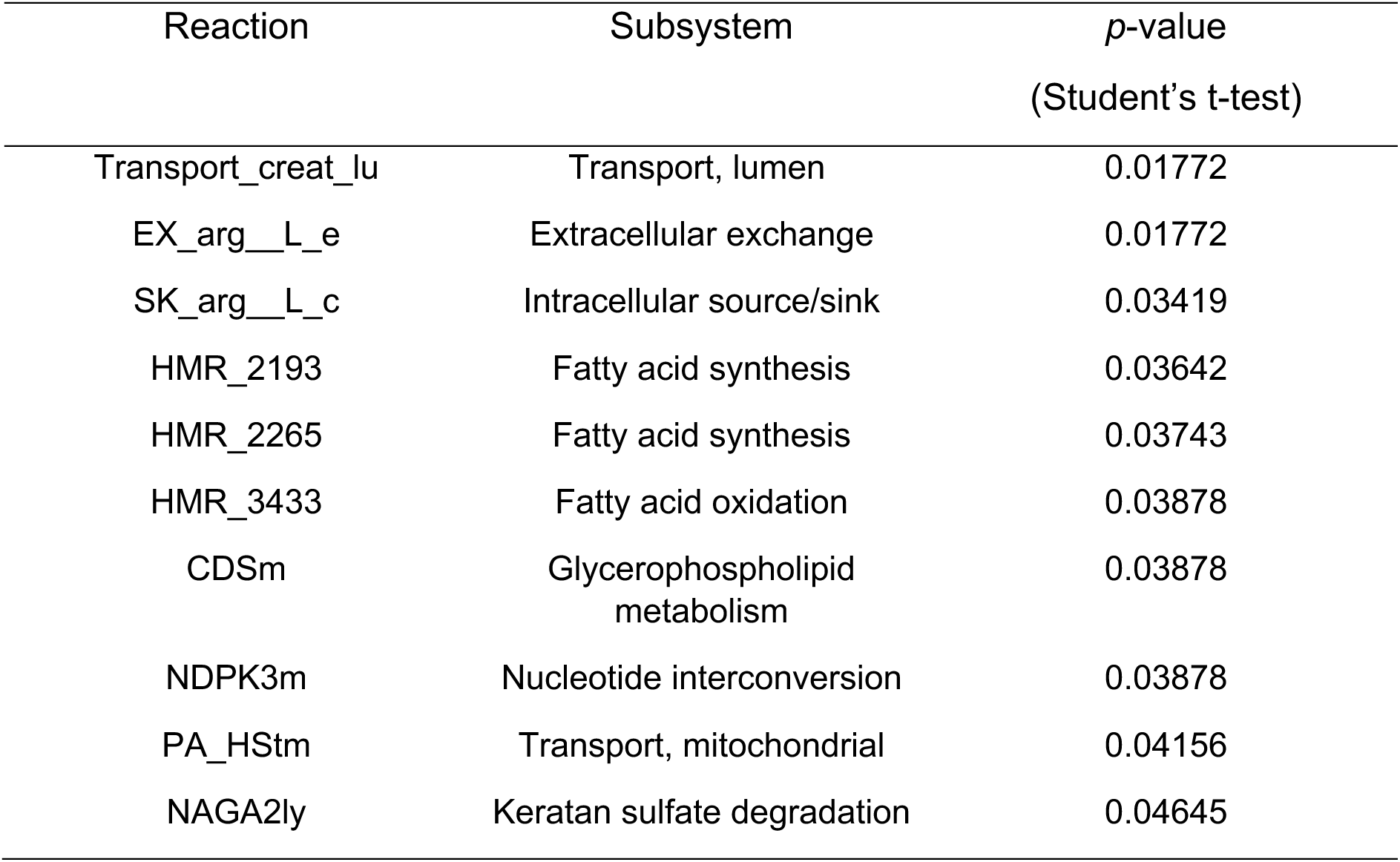
Comparison of reaction maximum fluxes between inflammatory and healthy groups in CD and UC, respectively. Table 3a: CD study: Different maximum fluxes between healthy and inflammatory groups

**Table 3b:** UC study: Different maximum fluxes between healthy and inflammatory groups

| Reaction | Subsystem | *p*-value<br>(Student's t-test) |
| --- | --- | --- |
| EX_dgchol_e | Extracellular exchange | 1.85113E-05 |
| UGALNACtg | Transport, golgi apparatus | 2.47163E-05 |
| SO4tl | Transport, lysosomal | 0.00096 |
| PAPtg | Transport, golgi apparatus | 0.00222 |
| NACHEX25ly | Keratan sulfate degradation | 0.00345 |
| IDPtn | Transport, nuclear | 0.00462 |
| DIDPtn | Transport, nuclear | 0.00462 |
| NDPK9n | Nucleotide interconversion | 0.00462 |
| NDPK10n | Nucleotide interconversion | 0.00462 |
| r0611 | Miscellaneous | 0.00462 |
| UDPtI | Transport, lysosomal | 0.00488 |
| GLCNACASE4ly | Heparan sulfate degradation | 0.00590 |
| ELAIDCRNt | Fatty acid oxidation | 0.00974 |
| UAGDP | Aminosugar metabolism | 0.01320 |
| LDH_D | Pyruvate metabolism | 0.01363 |
| ALAR | Alanine and aspartate metabolism | 0.01440 |
| RE3273C_1 | Phosphatidylinositol phosphate metabolism | 0.01578 |
| Transport_cmp_lu | Transport, lumen | 0.03249 |

Additionally, we carried out Monte Carlo Artificially Centered Hit and Run (ACHR) flux sampling to evaluate flux distributions belonging to tryptophan metabolism and nucleotide interconversion from the context-specific reconstructions. Flux sampling allowed us to perform an unbiased assessment of all possible flux distributions in the solution space of each context-specific reconstruction. For tryptophan metabolism, we identified that UC colonocytes showed lower flux distribution for the reaction that deaminates serotonin into 5-Hydroxyindoleacetaldehyde (5HOXINOXDA) (Figure 2A). CD colonocytes showed similar flux distributions for this particular reaction compared to healthy colonocytes. CD colonocytes showed higher fluxes in the conversion of acetoacetyl-CoA to crotonoyl-CoA (HACD1m and ECOAH1m), while these fluxes did not vary significantly between UC and healthy colonocytes. UC and healthy colonocytes had similar flux distributions for nucleotide interconversion among ATP, GDP, UDP, CDP and IDP, while CD colonocytes showed more variation in nucleotide interconversion when compared to healthy colonocytes (Figure 2B).

**Figure 2.**
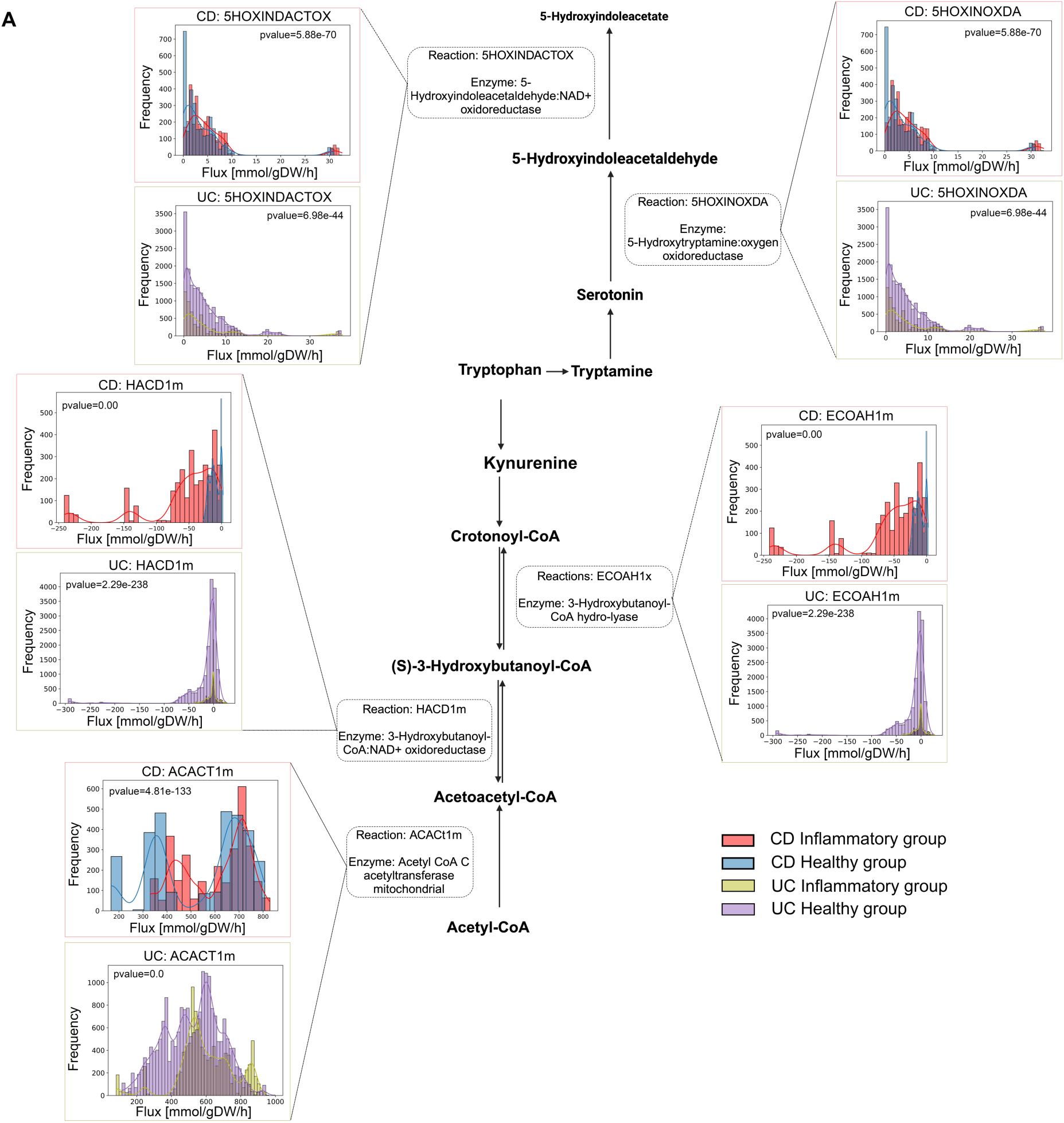
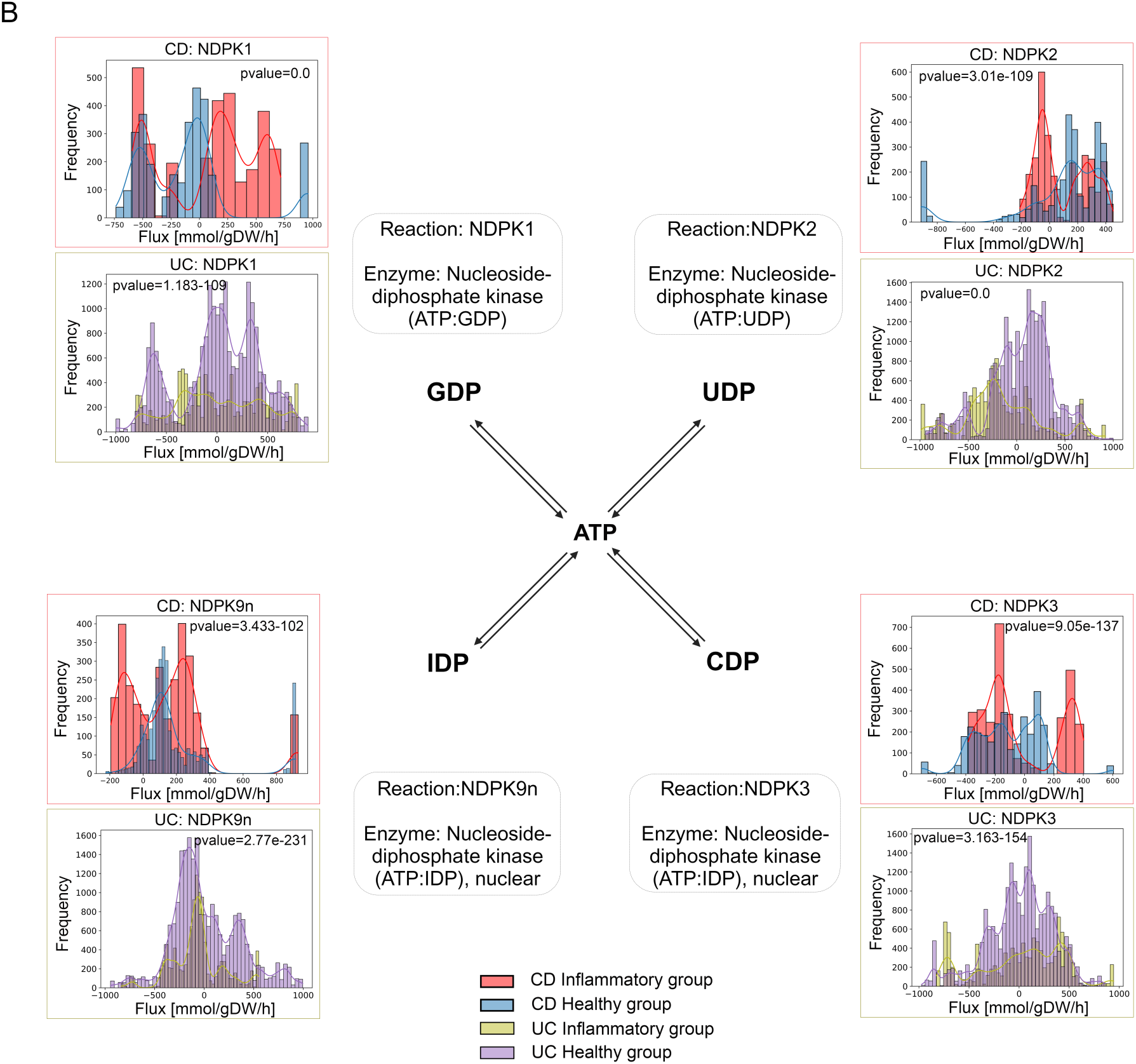
A) Bar plots indicating the comparison of flux distributions in tryptophan metabolism pathways between inflammatory and healthy group colonocytes from CD and UC, respectively. P-values were calculated with Kolmogorov-Smirnov test, reactions in the figure are the BIGG reaction ID; B) Bar plots indicating the comparison of flux distributions in nucleotide interconversion pathways between inflammatory and healthy group colonocytes from CD and UC, respectively. P-values were calculated with Kolmogorov-Smirnov test, reactions in the figure are the BIGG reaction ID.

#### In silico gene knockout simulations identify metabolic pathways affected in UC and CD

GEMs contain information on gene-protein-reaction (GPR) associations in the system that is useful for integration of multi-omics data. We can perform *in silico* single gene knockout to predict essential genes in the reconstruction as well as identify the effects of genes in the system. For the present case study, we performed *in silico* single-gene knockout simulation on all context-specific metabolic reconstructions in both CD and UC studies and compared the output of the biomass objective function. Changes in metabolic fluxes within biomass objective function could represent changes in cell viability (Brunk et al., 2018). The result of knockout simulations in CD colonocytes predicted that genes associated with glyoxylate and dicarboxylate metabolism (*GRHPR*), cholesterol metabolism (*HMGCS2*), CoA synthesis (*PPCS*), and transportation of metabolites (*SLC5A11* and *SLC5A7*) had significant effects on viability. In the UC colonocytes, single gene knockouts that significantly affected the biomass objective function were associated with carnitine metabolism (*CRAT*), glyoxylate and dicarboxylate metabolism (*ALDH9A1*) and transportation of metabolites (*SLC4A7*, *CRAT* and *CFTR*) (Figure 3b). This suggests the importance of several pathways in the context of IBD, like glyoxylate and dicarboxylate metabolism, as this pathway could affect objective functions of CD and UC reconstructions. Similar observations have also been reported in a study employing gut microbiome GEMs, which showed associations among glyoxylate and dicarboxylate metabolism, metabolic disorders and inflammation (Proffitt et al., 2022).

**Figure 3.**
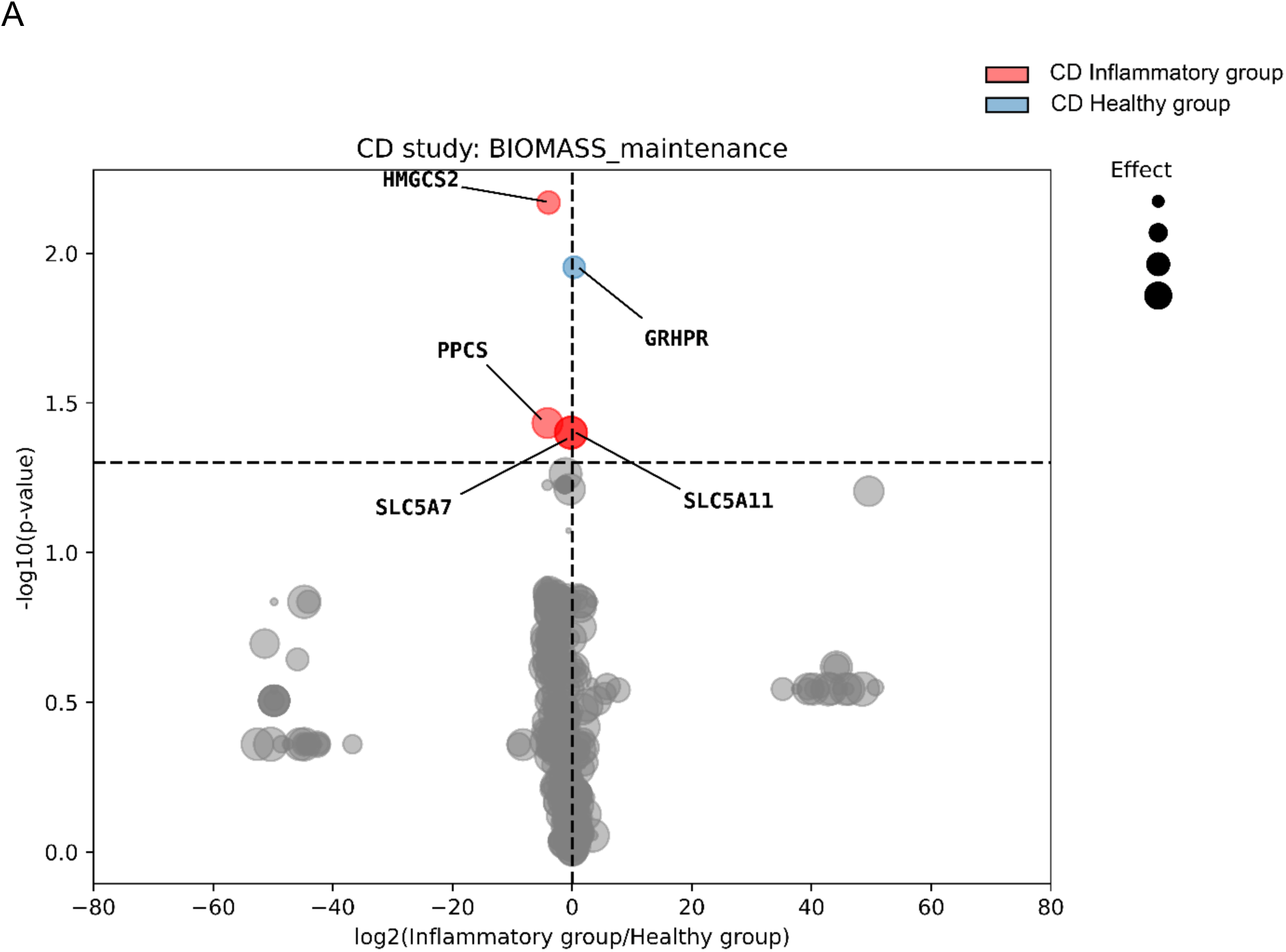
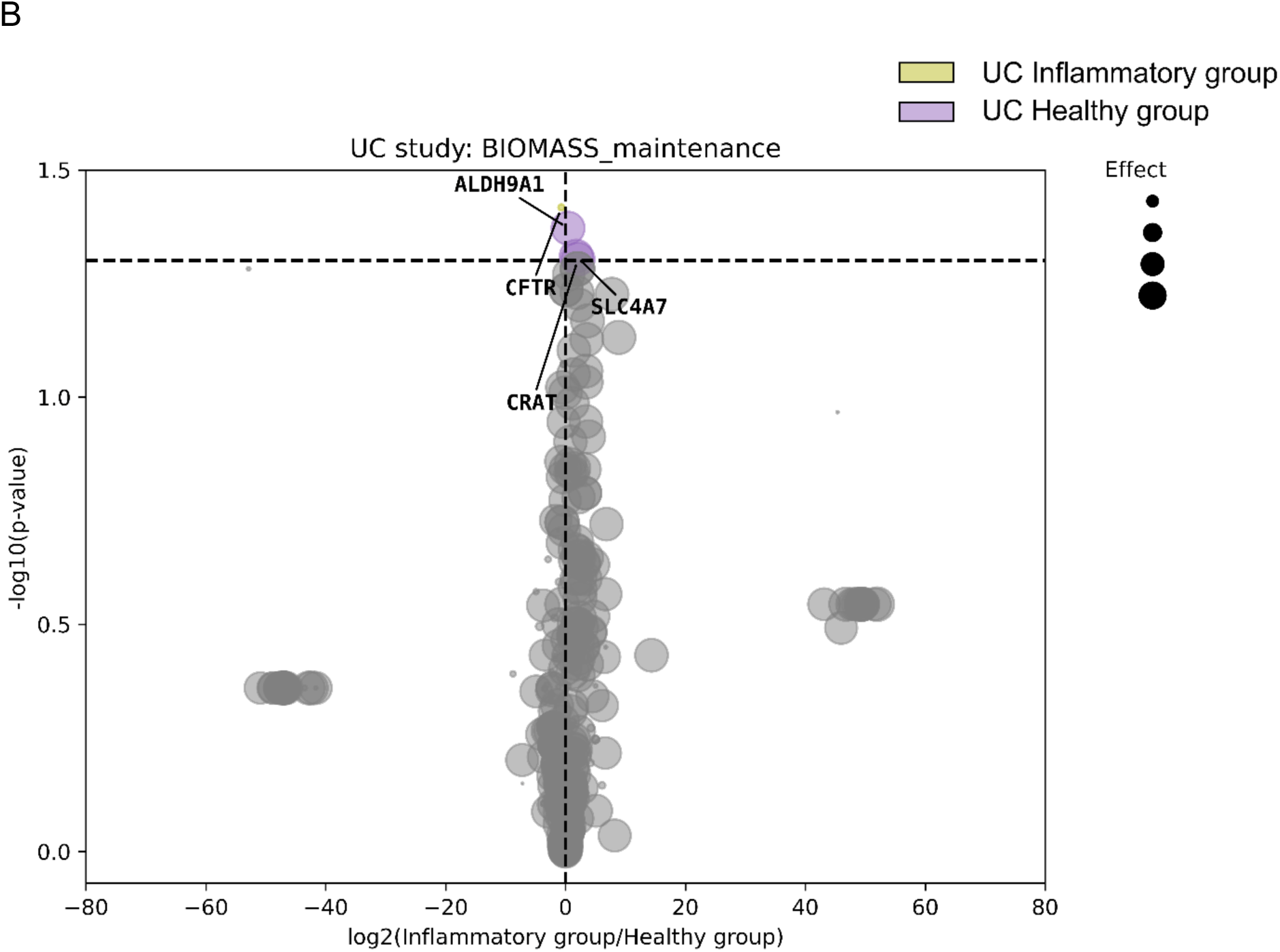
A) Volcano plot comparing effects of single gene knockout simulations on the colonocyte BIOMASS_maintenance flux between inflammatory and healthy group colonocytes from CD. X-axis: log_2_(Inflammatory group/Healthy group), where “Inflammatory group” represents the mean (WT-KO)/WT ratio in colonocytes of inflammatory group, and “Healthy group” equals the mean (WT-KO)/WT ratio in colonocytes of healthy group; values < 0 (colored dot on the left of the vertical dotted line) signify knockouts with greater effects on BIOMASS_maintenance in colonocytes of inflammatory group, whereas values > 0 (colored dot on the right of the vertical dotted line) signify knockouts with greater effects on BIOMASS_maintenance in colonocytes of healthy group. Y-axis: statistical significance (p-values calculated by student’s t-test) comparing knockout effects between inflammatory and healthy groups; dots above the horizontal dotted line (t-test p-value ≤ 0.05) are statistically significant. The size of each dot is proportional to the overall effect size (mean (WT-KO)/WT ratio across all inflammatory and healthy groups).; B) Comparison of effects of single gene knockout simulations on the colonocyte BIOMASS_maintenance flux between inflammatory and healthy group colonocytes from UC.

Comparing effects of gene knockouts between the IBD and healthy control groups showed that these genes played different roles in the maintenance of the biomass objective function. *PCYT2* gene knockouts were shown to cause a greater decrease in the objective function of CD colonocytes than healthy colonocytes. Knockouts of 5 genes (*PPCS*, *HMGCS2*, *GRHPR*, *SLC5A11* and *SLC5A7*) were predicted to cause a greater decrease in the objective function in CD colonocytes than healthy colonocytes. Compared to CD, we identified additional genes (*SLC4A7*, *CRAT* and *CFTR*) predicted to have a greater effect on decreasing flux of objective function in UC colonocytes. The gene knockout simulations reveal key metabolic pathways and transport mechanisms that significantly influence cell viability in IBD conditions. The differential effects of specific genes between CD and UC colonocytes, as well as between IBD and healthy controls, underscore the importance of pathway-specific interventions.

## Discussion

In this study, we present a metabolic reconstruction of human colonic epithelial cells, iColonEpithelium, which is useful for predicting metabolic phenotypes in the gut epithelium at the cellular level. The following are the key features of this cell-specific reconstruction: a) this is the first reported cell-type-specific metabolic reconstruction of human colonic epithelial cells built from transcriptomics data of clinical biopsies; b) colonic epithelial cells are known to interact with the gut microbiome, so we added a unique compartment named ‘lumen’ to enable simulation of metabolite exchanges between colonic epithelial cells and the gut microbiome; c) the iColonEpithelium reconstruction was able to simulate the transport of metabolites from the gut microbiome environment, as well as transport of metabolites into the portal vein; d) the reconstruction achieved cell-type specificity based on successful completion of several metabolic tasks important for colonic epithelial cells; e) through the integration of patient-derived single cell transcriptomic data with metabolic reconstruction, we identified metabolic alterations in ulcerative colitis (UC) and Crohn’s disease (CD), highlighting the utility of this *in silico* metabolic reconstruction.

The human colonic epithelium consists of multiple cell types, including colonocytes, goblet cells, enteroendocrine cells, tuft cells, and other immune cells. Compared to other cell types in the colonic epithelium, the colonocytes are the most abundant cell type and take a central role in nutrient exchanges with the gut microbiota and dietary substrates (Kong et al., 2018). Recent studies have highlighted the importance of colonocytes in modulating host-microbiota metabolic interactions. For instance, colonocytes are the tissue interface where SCFAs are absorbed and consumed by the host. The consumption of SCFAs, like butyrate and acetate, by colonocytes will affect the availability of these SCFAs for the rest of host body, and β-oxidation of SCFAs in colonocytes is accompanied by a high demand for oxygen, which has a direct effect on maintaining the anaerobic environment within the colon (Litvak et al., 2018b). However, heterogeneous uptakes of SCFAs by colonocytes under different conditions, like changed diet patterns or progression of gastrointestinal diseases, still remain unknown. Studies have indicated strong interactions among the gut microbiota, SCFA production, and colonocyte metabolism (Den Besten et al., 2013), A high-quality colonocyte-specific metabolic model has the potential to provide deeper insights into host-microbiome interactions in the gut.

To ensure the cell-type specificity of the iColonEpithelium reconstruction, the draft reconstruction was initially built with publicly available transcriptomics data of human colon epithelium biopsy samples, consisting largely of colonic epithelial cells. During the refinement and test of our draft reconstruction, we used metabolic tasks that have been reported for human colonic epithelial cells to validate that iColonEpithelium could carry out metabolic functions in accordance with reported experimental studies on the colonocytes. We also employed targeted metabolomics analysis of differentiated Caco-2 cells to determine the utilization of specific metabolites and used these results to refine the iColonEpithelium to make sure that it can capture metabolic functions specific to colonic epithelial cells.

In this study, we also integrated the scRNAseq data of colonocytes from two studies of CD and UC with iColonEpithelium to explore and compare metabolic changes in IBD. It has been reported that IBD is accompanied by metabolic dysfunctions in colonic epithelial cells (Rath & Haller, 2022). Our reconstruction could differentiate metabolic changes in CD and UC colonocytes, relative to healthy controls. For instance, in the CD, we predicted alterations in fatty acid metabolism, and recent studies have highlighted the importance of long chain fatty acids (LCFAs) in the CD (Piotrowska et al., 2021). For the UC, we saw disruptions in nucleotide metabolism, which were catalyzed by nucleoside-diphosphate kinase (*NDPK*) genes. The downregulation of *NDPK* in UC has also been reported previously (Crittenden et al., 2018). Both LCFAs and nucleotide metabolism were correlated with inflammation, which may provide insight into the different mechanisms underlying CD and UC.

We found different patterns in tryptophan metabolism between CD and UC. Tryptophan is an essential amino acid from the diet and its metabolism in the body is mediated by both the gut microbiota and colonic epithelial cells (S. Wang et al., 2023). According to the results (Figure 2A), CD colonocytes showed an increased flux distribution in reactions that degraded serotonin into 5-hydroxyindoleacetate (5-HIAA). It has been reported that the concentration of 5-HIAA from the urine samples of CD patients, which were collected in the afternoon, was higher than the afternoon urine from healthy individuals. (Bodriagina et al., 2019). We also identified a significant increase of nicotinamide adenine dinucleotide (NAD+) production from (S)-3-Hydroxybutanoyl-CoA (Figure 2A, reaction: HACD1m) in the colonocytes from CD patients when compared to colonocytes from healthy individuals. Tryptophan is an import precursor to NAD+ production, which serves as a cofactor and participates in a wide range of homeostatic processes. In IBD, dysregulated NAD homeostasis will not only compromise the intestinal barrier but also perturb mucosal immunity (Xue et al., 2023). From the analysis in this study, we also highlight the importance of metabolic flux in HACD1m in CD colonocytes.

Single gene knockout simulation allowed us to explore essentiality of genes between CD and UC colonocytes, and some of the predicted essential genes are in accordance with experimental results. For example, a study of *in vitro* model of CD reported upregulated expression levels of *GRHPR*, and knock-down of *GRHPR* could increase apoptosis of intestinal epithelial cells (Zong et al., 2016). This validation further substantiates the capacity of our reconstruction to capture important metabolic processes in colonocytes, when combined with -omics data.

In summary, we constructed the first cell-type-specific GEM of human colonic epithelial cells to simulate and explore their metabolic activities. The iColonEpithelium has the capacity to achieve essential metabolic functions of human colonic epithelial cells. When constrained with scRNAseq data from IBD patients, it could predict and capture some important metabolic changes in CD and UC, like fatty acid metabolism tryptophan metabolism, which were in accordance with experimental results reported in literature. The iColonEpithelium reconstruction can contribute to our understanding of metabolism in colon epithelium, and we have the ability to combine our model with gut microbiota GEMs, through the integration of a luminal compartment, to further explore effects of dietary and microbiome variation on host metabolism.

## Limitations of the study

We recognize that our work has certain limitations. Our *in silico* metabolic reconstruction was built from existing transcriptomic data of human colon epithelium, which contains comprehensive information of genes expressed in a given cell type. We need more data to add other constraints, like enzyme kinetics and thermodynamic parameters. Our current predictions are all assuming the steady state condition. Dynamic simulation requires the estimation of unknown parameters, like the rate constants of reactions, and the validation of these estimated parameters requires time-course experimental measurements (Borzou et al., 2023). Unfortunately, these measurements are not available for human colonocytes. For the case studies, we were able to obtain only scRNAseq data from CD and UC samples. There are no human colonocyte studies with matched metabolomics, proteomics and transcriptomics data, which could be used to further constrain our models. Also, there is limited information on the metabolites in the extracellular space available to the colonocytes. Future work should integrate proteomics data from colonocytes to further constraint the GPR parameters. Availability of metabolomics data from both gut microbiota and blood can be helpful in constraining the reconstructions and validate the predictions.

## Methods

### Collection of transcriptome data and cross-platform normalization

We searched NCBI’s GEO database (https://www.ncbi.nlm.nih.gov/geo/) and collected transcriptome data of human colonic epithelial cell from published studies with the following criteria: (1) clear description of sample sources, which were from human colon epithelium; (2) inclusion of samples from healthy volunteers; (3) availability of raw data; (4) data published within the last 10 years. As we collected both RNA-seq and microarray data from various platforms based on the above-mentioned criteria, we further applied a recently published tool – Shambala-2 (Borisov et al., 2022) to perform cross-platform normalization. According to the workflow of Shambala-2, we normalized all RNA-seq and microarray raw data separately and then combined them together. The datasets included in present study are described in details in Table S2.

### Preparations of inputs for different GEM reconstruction tools and pruning draft reconstructions

We used four prominent reconstruction extraction tools to obtain draft reconstructions for colonocyte, using Recon3D (Brunk et al., 2018) as the template reconstruction. Each of them uses a different algorithm and has different requirements for inputs. The output draft reconstructions from each method were compared and analyzed to ensure comprehensiveness and specificity of our colonic epithelial cell reconstruction. The four methods for generating cell-specific reconstruction are described below:

*pymCADRE*: pymCADRE (v1.2.3, https://github.com/draeger-lab/pymCADRE/) is the python version of the mCADRE (Leonidou et al., 2023).We first calculated the ubiquity score of genes. After comparing different quantiles of gene values, we used the 30th percentile expression value as the cutoff for gene expression. Then all gene values are binarized according to the cutoff to calculate a ubiquity score of each gene (Y. Wang et al., 2012). Then, we prepared the precursorMets list which included all metabolites involved in the ‘BIOMASS_maintenance’ reaction and the confidence score of all reactions in Recon3D. We obtained the draft reconstruction using these input parameters and employing the pymCADRE algorithm.

*iMAT* and *tINIT*: python version of both iMAT and tINIT are integrated into the troppo (v0.0.7, https://github.com/BioSystemsUM/troppo). Similar to the pymCADRE protocol, we set the 30th quantile of normalized expression data as our gene expression cutoff. Then we took median values among all samples as gene expression values. All values below the cutoff were converted to 0, while values over the cutoff were log2 transformed. Using the gene expression values, we obtained the reaction scores in troppo (Ferreira et al., 2020).

*CORDA*: CORDA (v0.17, https://github.com/resendislab/corda) is also a reconstruction tool in python. We used the same reaction scores as iMAT and tINIT and recalculated them at three expression levels, as required in CORDA (Schultz & Qutub, 2016). The target metabolites required by CORDA were the same as those on the pymCADRE precursorMets list.

We used pymCADRE, iMAT, tINIT and CORDA to get four draft reconstructions, that were used for generating the consensus reconstruction.

### Generation of consensus reconstruction

We identified overlapping reactions, genes and metabolites for the draft reconstructions, and built a consensus reconstruction which included all reactions from the four draft reconstructions except reactions unique to the draft reconstructions of tINIT. The draft reconstruction generated by tINIT contained the most reactions. After running FASTCC, the output of tINIT draft reconstruction contained the greatest number of unique metabolites and genes among the four reconstructions, implying a higher level of uniqueness of the tINIT reconstruction. Therefore, we excluded reactions unique to tINIT and merged other reactions in the four draft reconstructions to build a consensus reconstruction for the following analyses.

### Metabolic task analysis for specificity testing

We tested the specificity of our consensus reconstruction using the information from 229 reported metabolic functions known to be feasible for human colonocytes as well as general human cells (supplementary Table S4). In brief, the reconstruction was closed by setting lower boundaries of both exchange and sink reactions to 0. Then for each metabolic task, reactants would be available for the reconstruction by setting the corresponding exchange reactions or sink reactions to -1. We added a sink reaction for the product of each metabolic task and set it as the objective. We ran model.optimize function in COBRApy (Ebrahim et al., 2013) to optimize the sink reaction to check if it could achieve a flux value over 0.

### Curating the transport reactions module

To allow our human colonic epithelial cell reconstruction to connect and interact with the gut microbiome, we built a lumen compartment containing transport reactions which enable the exchange of metabolites between colonic epithelial cells and the gut microbiome. We collected genes associated with transport functions from the Human Protein Atlas (www.proteinatlas.org) (Uhlen et al., 2019) and also retained the genes which were part of the consensus reconstruction (Supplementary Table S5). Then, we searched metabolic reactions related to these genes in Uniprot (Bateman et al., 2023), after which metabolites from these reactions were extracted. Meanwhile, we also extract metabolites from a published human membrane transporter module (Sahoo et al., 2014), and colon transport reactions of the human whole-body model (Thiele et al., 2020). After curating a list of metabolites from sources described above, we identified those that overlapped with the consensus reconstruction. Finally, transport reactions associated with these metabolites were added to the lumen compartment to allow metabolite exchanges with gut microbiota.

### Testing basic properties of reconstruction

The quality of our reconstruction was checked with a series of tests implemented in the COBRA Toolbox (Heirendt et al., 2019), including leak tests and sanity checks (Thiele et al., 2020). During the tests, all demand, sink and exchange reactions in our reconstruction were closed, and then the closed reconstruction was checked for (1) whether it could produce any exchange metabolites with or without a demand reaction for each metabolite; (2) whether it could produce energy from water or oxygen; (3) whether it could produce any metabolites with a reversed ATP demand reaction; (4) whether it could achieve flux consistency – dead ends in the reconstruction were detected and excluded with FVA (Thiele et al., 2013).

### Differentiated Caco-2 cell lines and metabolomics analysis

Caco-2 cells were purchased from American Type Culture Collection (Manassas, VA) and cultured in a 37°C humidified incubator with 5% CO2 utilizing T-75 flasks (FB012937, Fisherbrand), with Dulbecco’s Modified Eagle Medium (DMEM) (10-013-CV, Corning, supplemented with 10% fetal bovine serum (FBS) (S11150, R&D). To induce differentiation, the medium was replaced every 2-3 days. Cells were passaged using trypsin (25-03-CI, Corning). At the end of the experimental timeline, culturing medium of the Caco-2 cells and fresh DMEM + FBS was collected for targeted metabolomics analysis of SCFAs (butyrate, acetate and propionate) and 20 aminoacids (Table 2). The concentration of SCFAs in samples was detected using a reported liquid chromatography with tandem mass spectrometry (LC-MS-MS) method (Chan et al., 2017) in the Bindley Bioscience Center at Purdue University.

### Analysis of IBD patient-derived samples

We collected scRNAseq data of human colon epithelium from two studies: a CD study (Kanke et al., 2022) and a UC study (Smillie et al., 2019). They both contained more than three samples each in the IBD groups and healthy control groups. Colonocyte gene expression matrices from IBD samples and healthy samples, for both CD and UC studies, were extracted using the Seurat package in R (Hao et al., 2021). Then, using iMAT, we integrated gene expression data of each sample with our iColonEpithelium to build context-specific reconstructions of all samples. We ran FVA on each context-specific reconstruction and performed student’s t-test (alpha < 0.05) on the maximum fluxes between IBD and healthy control groups. This analysis led to the identification of fluxes with significant differences across UC and CD vs healthy control groups. Genes associated with these differential reactions were also collected.

In addition to FVA, we also employed the ACHR flux sampling function in the COBRApy with a thinning factor of 1000. This provided us with random samples of steady-state fluxes with the allowable solution space of each context-specific reconstruction. Solutions from flux sampling were used to compare the flux distributions between IBD and healthy control groups.

We performed single gene deletion simulation on all context-specific reconstructions. In brief, we knocked out one gene at a time, and then ran FVA to obtain the maximum flux of specific reactions. Then according to a method from a published study (Lewis et al., 2021), we calculated the ratio of altered maximum fluxes caused by a single gene knockout in the IBD and healthy control groups, respectively. Finally, ratios between the IBD and healthy groups were compared.

## Acknowledgements

Research reported in this publication was supported by the National Institute of Diabetes and Digestive and Kidney Diseases of the National Institutes of Health under award number R01DK133468 (to S.M.G.).

## Author information Contributions

B.J. generated the metabolic reconstructions, searched for experimental data, performed simulations and interpreted the data. B.J, N.Q-B, C.D, S.M.G and P.B. contributed to interpretation of results. V.B.N. and L.R. performed the experiments on differentiated Caco-2 cells. Y.H-H. performed metabolomics analysis and B.J. and Y.H-H. interpreted the metabolomics results. P.B. and S.M.G. conceptualized, supervised and provided resources. B.J. and P.B. drafted the manuscript. All authors participated in editing and revising the manuscript.

## Data availability

### Code

Jiang, B. (2024), “IColonEpithelium_GEM.” (DOI: 10.4231/FD22-9D96).

